# Influence of Hypoxia on a Biomaterial Model of the Bone Marrow Perivascular Niche

**DOI:** 10.1101/2025.02.20.639296

**Authors:** Gunnar B. Thompson, Victoria R. Barnhouse, Sydney K. Bierman, Brendan A.C. Harley

## Abstract

Hematopoietic stem cell (HSC) fate is shaped by distinct microenvironments termed niches within the bone marrow. Quiescence, expansion, and differentiation are directly and indirectly regulated by complex combinations of cell secretomes, cell-cell interactions, mechanical signals, and metabolic factors including oxygen tension. The perivascular environment in the bone marrow has been implicated in guiding HSC fate. However, bone marrow presents an environment which is hypoxic (∼1-4% O_2_) relative to traditional cell culture conditions, and the study of hypoxia in vitro is complicated by the speed with which normoxic conditions during HSC isolation induce differentiation. There is a unique opportunity to use engineered models of the bone marrow to investigate the impact of defined hypoxia on HSC fate. Here, we examine the coordinated impact of oxygen tension and the perivascular secretome upon murine hematopoietic stem and progenitor cells (HSPCs) in vitro. Our findings highlight the importance of mitigating oxygen shock during cell isolation in engineered marrow models. We report a shift toward the Lineage^-^ phenotype with hypoxic culture, expansion of HSPCs in response to perivascular niche conditioned medium, and enhanced HSPC maintenance in a hydrogel model of bone marrow in hypoxic culture when oxygen shock is mitigated during isolation using cyclosporin A.

## 1. Introduction

Hematopoietic stem cells (HSCs) reside in the bone marrow and through the process of hematopoiesis, produce the full complement of blood and immune cells needed throughout a person’s lifetime^1^. They are rare within the bone marrow and are prospectively identified using flow cytometry. Most commonly, a compartment of hematopoietic stem and progenitor cells (HSPCs) is identified as Lin^-^Sca-1^+^c-Kit^+^, and this population is typically referred to as the LSK population^2,3^. This compartment further comprises long-term repopulating HSCs, short-term repopulating HSCs, and multipotent progenitors (LT-HSCs, ST-HSCs, and MPPs respectively) which may be phenotypically identified by their expression of CD150 and CD48, and are functionally distinct in their self-renewal and repopulating capacity^4,5^. Though not fully understood, it appears that more stemlike HSCs, such as the CD150^+^CD48^-^CD41^-^ subfraction are associated with greater levels of quiescence^4,6^. HSC transplants are used to cure diseases including acute myeloid leukemia, sickle cell disease, and numerous others^7^. Unfortunately, a significant number of post-transplant relapses occur when donor cells fail to engraft, which is often due to a low cell dose^8,9^. One possible avenue to mitigate this risk is pre-transplant expansion of stem cells to increase the delivered cell dose. While significant strides have recently been made on this front^10,11^, expansion remains a challenge in the field of hematology which relies heavily upon conventional 2D cell culture. A better understanding of how HSCs interact with and are positively regulated by their native bone marrow microenvironment may eventually provide the scientific leverage necessary for clinical expansion of HSCs.

HSCs have been identified in several unique locales of the bone marrow, termed niches, which provide a set of matrix, cellular, and biomolecular signals that influence HSC growth, quiescence, and lineage specification^12^. The two most frequently named niches that differentially support HSCs are the endosteal niche, which is proximal to the inner surface of the bone, and the perivascular niche (PVN) which is sometimes further divided into the periarteriolar and perisinusoidal niches^13,14^. The bone marrow is an overall hypoxic tissue, with oxygen levels varying between 1-4% depending on location^15^, and with the highest oxygen tension near the surface of bone and arterioles, and the lowest toward the middle of the marrow and near sinusoids^15,16^. Niche hypoxia is believed to play an important role in HSC activity by helping to maintain a fraction of quiescent HSCs via HIF-1α stabilization^17,18^. Work published by the Broxmeyer Lab demonstrated the importance of oxygen tension during cell culture and during cell isolation, finding that isolation of HSCs in ambient air with high oxygen concentrations quickly leads to extraphysiologic oxygen shock/stress (EPHOSS)^19–22^. Here, normoxic isolation leads to increased reactive oxygen species (ROS) generation and mitochondrial activity, mitochondrial permeability transition pore (MPTP) induction, and HSC differentiation. However, this phenotype could be mitigated via either complicated all-hypoxic isolation or via using cyclosporin A (CSA) to inhibit cyclophilin D, a key factor in MPTP induction. These studies showed that hypoxic (3% O_2_) or CSA-supplemented HSC isolation from the bone marrow inhibited differentiation and enhanced transplant efficacy in mice.

The vast majority of in vitro hematopoietic niche model have been performed in a conventional incubator with ∼18% O_2_^23^, which greatly exceeds physiological oxygen levels in bone marrow. One method of studying the effect of low oxygen levels is to use a chemical inducer of hypoxia^24^. Braham et al. used cobalt (II) chloride hexahydrate to induce a hypoxic environment and saw a decrease in cell proliferation^25^ but did not evaluate phenotypic markers of HSPC response. Others used oxygen-controlled (hypoxic) incubators: for example, Amiri et al. co-cultured human HSCs with Wharton jelly mesenchymal stem cells (WJ-MSCs) in hypoxia vs. normoxia^26^. Interestingly, WJ-MSC spheres cocultured with HSCs in hypoxia increased numbers of HSCs and long-term culture initiating cells, though the effect appears primarily driven by WJ-MSC spheres rather than hypoxia. Araki et al. showed hypoxic culture reduced ROS and enabled greater human HSPC expansion via Delta-like ligand 1 and Notch signaling^27^. Finally, Igarashi et al. showed selective expansion of HSCs in 2D hypoxia cultures^28^. These findings motivate efforts to create biomaterial models of the bone marrow to examine the combination of hypoxia and the niche on HSPC expansion.

We have previously developed a bone marrow PVN model for culture of murine HSPCs within a methacrylated gelatin (GelMA) hydrogel^29,30^. These studies examined the role of both PVN conditioned media and direct co-culture with an engineered PVN, finding PVN-derived angiocrine factors promoted both the retention of a subset of HSPCs as well as expansion of terminally differentiated hematopoietic cell populations. Here, we examine the influence of extracellular hypoxia and PVN-derived factors on HSPC activity. We first examine how exposure to continuous hypoxia (1.5% O_2_) alters the morphology of the endothelial networks as well as their secretome. We then analyze the response of HSPCs to hypoxia within the three-dimensional hydrogel in the presence or absence of conditioned media from the engineered PVN model. Finally, we use CSA to mitigate the effects of EPHOSS during HSC isolation and investigate the impact of sustained hypoxic exposure and the perivascular secretome on HSPC fate in a GelMA model of the bone marrow. These efforts seek to define the importance of EPHOSS-mediation as well as the role of metabolic and biomolecular signals from the marrow vascular microenvironment during in vitro studies of HSC expansion.

## 2. Materials and Methods

### 2.1. Cell Culture

Primary murine bone marrow derived endothelial cells (BMECs) were purchased from Cell Biologics (Chicago, IL) and murine mesenchymal stem cells (MSCs) from Cyagen (Santa Clara, CA). BMECs were cultured in complete mouse endothelial cell medium consisting of 5% FBS, EGF, hydrocortisone, VEGF, ECGS, heparin, l-glutamine and antibiotic-antimycotic solution (Cell Biologics). MSCs were cultured in mouse MSC growth medium with 10% FBS (Cyagen). Cells were cultured in 5% CO_2_ at 37° C. BMECs were used at passage 7 and MSCs were used at passage 10 in accordance with suppliers’ instructions.

### 2.2. Gelatin Methacrylate hydrogel formation and characterization

Type A Gelatin with 300 Bloom Strength (Sigma-Aldrich, St. Louis, MO) was functionalized with methacrylamide groups as described previously^29,31^. Briefly, methacrylic anhydride was added dropwise to a solution of gelatin and carbonate-bicarbonate buffer at a pH of 9-10^32^. After an hour, the reaction was stopped by adding warm deionized (DI) water. The pH was checked to ensure it was between 6-7, then the gelatin solution was dialyzed against DI water for one week and lyophilized. The gelatin:MA ratio was altered to control the degree of functionalization (DOF) and quantified by ^1^H NMR as previously described^31^. For this study, GelMA with a DOF of 63% relative to the phenylalanine peak was used for all experiments (**Fig S1A**). Hydrogels were formed by dissolving GelMA at 5wt% in PBS at 65° C, with 0.1% w/v lithium acylphosphinate (LAP) as photoinitiator. The prepolymer solution was pipetted into circular Teflon molds with a diameter of 5 mm and 1 mm thickness for cell culture experiments, then exposed to UV light (λ = 365 nm, 7.1 mW cm^−2^) for 30 s.

The stiffness of hydrogels was measured with unconfined compression on an Instron 5943 Universal Testing System (Instron, Norwood, MA) as previously described^33^. Briefly, hydrogels were compressed at a rate of 0.1 mm/minute and moduli were calculated using a custom MATLAB code^34^. Moduli were estimated from a fit to the linear viscoelastic regime of 0-15% strain beyond an initial offset of 2.5% above the noise floor^35^.

### 2.3 Endothelial Network Formation in Hypoxia or Normoxia

Endothelial networks were formed by co-culturing BMECs and MSCs in a 2:1 ratio (BMEC:MSC) with a total cell density of 2 x 10^6^ cells/mL. Cells were resuspended in the prepolymer solution, pipetted into Teflon molds, and photopolymerized as described above. Hydrogels were cultured for 7 days in a 1:1 blend of endothelial media and StemSpan SFEM HSC media (Stemcell Technologies, Vancouver, BC, Canada) with additional 50 ng/mL VEGF with daily partial (2/3 volume) media changes, leaving ∼250 µL of media to allow cell secreted factors to remain in the culture. Media was equilibrated overnight in either the InVivO2 physiological workstation (Baker Ruskinn, Sanford, ME) or a conventional cell culture incubator to equilibrate with appropriate oxygen levels. Hypoxia gels were made at the same time as normoxia gels, then moved to the hypoxic incubator set to 1.5% O_2_, 5% CO_2_, and 37° C.

### 2.4 Immunofluorescent Staining of Endothelial Cell Networks

Hydrogels were fixed in formalin (Sigma Aldrich) and washed with PBS. Gels were then blocked in PBS with 2% bovine serum albumin (BSA, Sigma Aldrich) and 0.1% Tween 20 (Thermo Fisher Scientific, Waltham, MA). Cells were stained with primary antibodies overnight at 4 °C (**Table S1**), then washed and incubated with secondary antibodies overnight at 4 °C as previously described^30,36,37^. Gels were then soaked in PBS with 0.5% Tween 20 to permeabilize the encapsulated cells. Hoechst 33342 (Invitrogen) was added as a counterstain at 1:2000 in PBS for 30 min after the secondary staining. PBS with 0.1% Tween-20 was used to wash at all steps after permeabilization.

### 2.5 Image Acquisition and Analysis

Images of endothelial networks used for analysis of morphological characteristics were acquired with a DMi8 Yokogawa W1 spinning disk confocal microscope outfitted with a Hamamatsu EM-CCD digital camera (Leica Microsystems, Buffalo Grove, IL). Z-stacks were acquired with a depth of 200 μm and step size of 5 μm. Six regions were imaged per gel, and three gels were analyzed per condition. The networks were quantified for metrics of total network length, and number of branches and junctions, using a previously established computational pipeline implemented in ImageJ and MATLAB^30,38,39^. Briefly, a custom plugin in ImageJ filters and binarizes the z-stacks so that the vessels are extracted from the image. In MATLAB, the vessels are skeletonized and then converted into a quantifiable nodal network. As this approach generates a 3D skeleton of all vessels within an experimental volume, it is able to accurately resolve both the number of branches and vessel segments^40^. Protein deposition was visualized using either a Zeiss LSM710 multiphoton confocal microscope or a Zeiss LSM880 Airyscan confocal microscope. For analysis of overlap and areal coverage, z-stacks were compressed into a maximum intensity projection prior to analysis.

### 2.6 Cytokine Array

Conditioned media was collected on day 7 from both the hypoxia and normoxia endothelial network conditions. The Proteome Profiler Mouse Angiogenesis Cytokine Array (R&D Systems, Minneapolis, MN) was used to identify what the endothelial network co-cultures secreted. Each array was incubated with 1 mL of conditioned media, and membranes were imaged with an ImageQuant 800 (Cytiva, Marlborough, MA) using chemiluminescence and an exposure time of 4 minutes. Images were analyzed using the microarray profile plugin in ImageJ to measure the integrated density of the dots. The measurements were background-subtracted, then normalized to the positive control spots.

### 2.7 Normoxic, CSA-supplemented, and Hypoxic Hematopoietic Stem and Progenitor Cell Isolation

All work that utilized primary cells was conducted under approved animal welfare protocols (Institutional Animal Care and Use Committee Protocol #23033, University of Illinois Urbana-Champaign). Murine LSKs were isolated from 4-8 week-old female C57BL/6 mice (The Jackson Laboratory, Bar Harbor, ME) as previously described^33^. Briefly, tibiae and femora were removed and crushed using a pestle and mortar to free cells from the marrow. The cell mixture was enriched first using a 1X red blood cell lysis buffer (Biolegend, San Diego, CA) followed by hematopoietic lineage negative enrichment using the EasySep^TM^ Mouse Hematopoietic Progenitor Cell Isolation Kit Stemcell Technologies). Cells were then stained for lineage markers, Sca-1, and c-Kit (**Table S2**) and sorted via florescence activated cell sorting using a Bigfoot Spectral Cell Sorter (Invitrogen).

### 2.8 Hematopoietic Stem Cell Culture in Hypoxia vs Normoxia

For studies of the effect of hypoxia and PVN conditioned media on stem cells, LSKs were encapsulated in GelMA hydrogels as described previously^41^, with 4500-6000 LSKs/gel. Hydrogels were maintained in 48-well plates for 4 days with media changes on day 2. Media was either a 50:50 blend of conditioned media collected from PVN hydrogel cultures and Stemspan SFEM (Stemcell Technologies) with a final concentration of 100ng/mL SCF (Peprotech, Cranbury, NJ) and 0.1% penicillin/streptomycin (PS, Gibco, Waltham, MA), or a control media of Stemspan SFEM with 100 ng/mL SCF and 0.1% penicillin-streptomycin. Gels were then cultured either in a normal cell culture incubator with ambient oxygen (∼18%) and 5% CO_2_, or an InVivO_2_ physiological workstation with O_2_ levels of 1.5% and CO_2_ at 5%.

On day 4, hydrogels were dissolved using 100 units collagenase type IV (Worthington Biochemical Corporation, Lakewood, NJ) on a shaker in the incubator at 37°C for 20 min, with gentle mixing by pipetting. Collagenase was quenched with 1 mL PBS + 5% FBS and centrifuged at 300 rcf x 5 minutes. The collected pellet was resuspended in PBS + 5% FBS and stained with a cocktail of antibodies (**Table S3**) to differentiate long term repopulating HSCs (LT-HSCs: CD150^+^CD48^-^LSK) and short term repopulating HSCs (ST-HSCs: CD150^-^CD48^-^ LSK), and multipotent progenitors (MPPs: CD150^+/-^CD48^+^LSK)^42,43^. Following staining, the cells were fixed and permeabilized with Foxp3/Transcription Factor Staining Buffer Set (Invitrogen). Cells were resuspended in PBS + 5% FBS and analyzed using a FACSymphony A1 flow cytometer (BD Biosciences, Franklin Lakes, NJ). Fluorescence minus one (FMO) controls were created by isolating Lin^-^ enriched bone marrow cells according to the primary cell isolation protocol described above.

### 2.9 Statistics

Normality of the data was determined using the Shapiro-Wilkes test^44,45^, and homoscedasticity using the Brown-Forsythe test^46,47^ at the 0.05 significance level. For experiments with two groups, a Student’s two sample t test was performed, or alternatively a Mann Whitney U test was used as a nonparametric alternative if normality and homoscedasticity were not satisfied. For 2 x 2 factorial experiments, a 2-way ANOVA test was used if normality and homoscedasticity were satisfied, followed with Tukey’s Honest Significant Differences test to compare groups. When those conditions were not satisfied, a Kruskal-Wallis test was used to evaluate main effects and a Pairwise Wilcoxon Rank Sum test was used to compare groups. The Benjamini & Hochberg correction was applied for statistical tests comparing multiple groups^48^. Number of replicates (n values) are reported in the figure captions. Graphs were made in R, and bar graphs represent the mean +/- the standard error of the mean (SEM) unless otherwise noted.

## 3. Results

### 3.1 Formation of Primary Endothelial Networks in Hypoxia and Normoxia

We first examined the ability of murine BMECs and MSCs to form endothelial networks under standard normoxic and hypoxic conditions: ∼18%^23^ and 1.5% oxygen, respectively. All hydrogels used in this study possessed a stiffness of 2.64 ± 0.28 kPa (**Fig S1B**), consistent with the bone marrow (0.25 – 24.7 kPa^49^). Maximum intensity projections for CD31^+^ cells reveal PVN networks in both normoxia and hypoxia (**Fig 1A**). While hypoxic culture induced networks with lower total length, less vessels, and less branch points (**Fig 1B-D, Table S4**) at day 7, the average branch length was similar for normoxic and hypoxic culture, suggesting similar vessel building blocks regardless of oxygen tension (**Fig 1E**). Not surprisingly, the composite metabolic activity of the endothelial networks showed a significant (p < 0.001) 28% reduction under hypoxic vs. normoxic conditions (**Fig 1F**).

**Figure 1.**
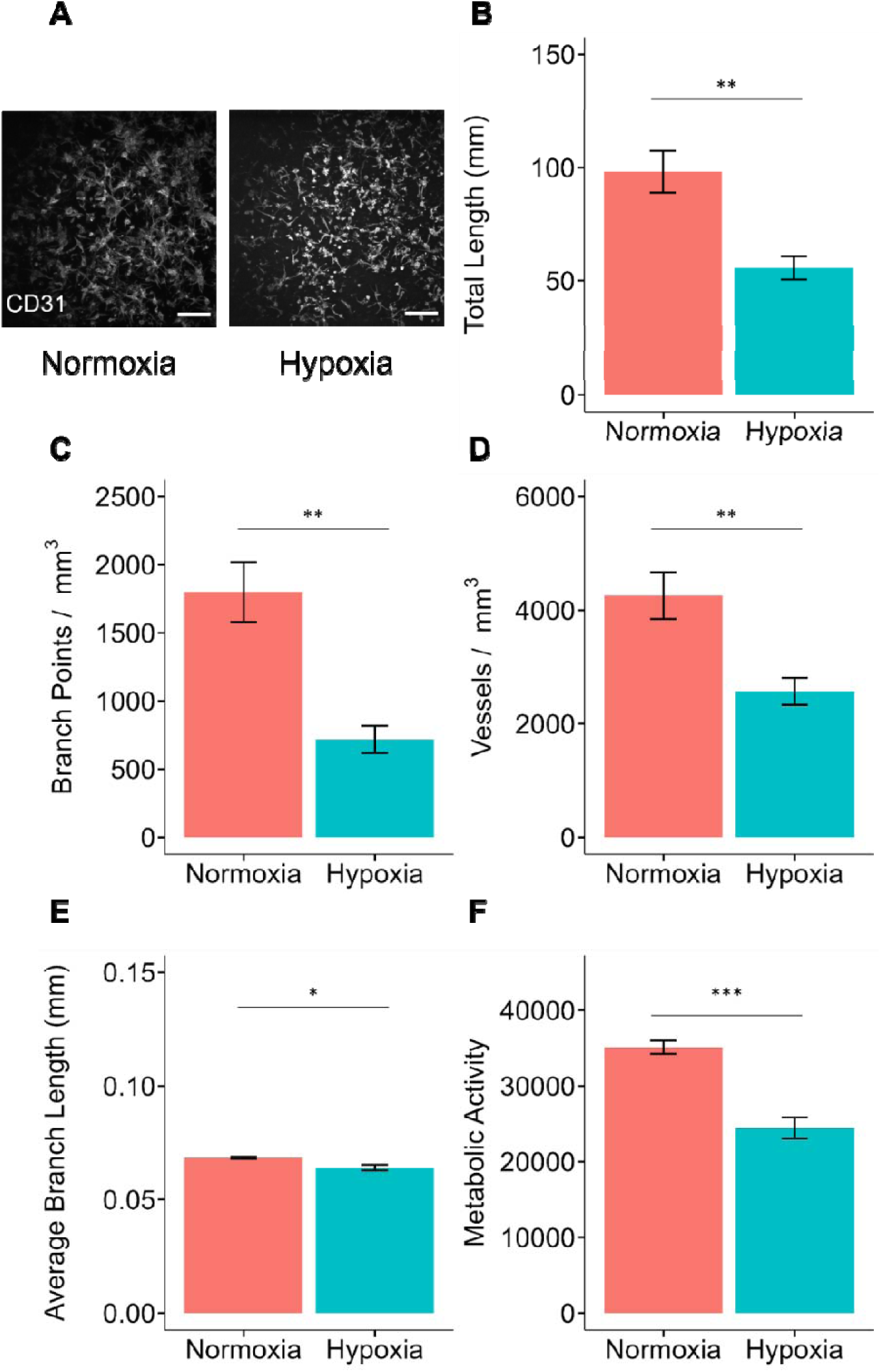
**A** CD31 staining reveals network formation in murine EC-MSC co-cultures grown in hypoxia vs. normoxia. Barplots present **B** total network length, **C** branch point density, **D** network density, and **E** average branch length for hypoxia- and normoxia-cultured networks. N = 6 gels for morphology studies. **F** Metabolic activity was assessed using alamarBlue assay. N = 9 gels. * indicates p < 0.05; ** indicates p < 0.01; *** indicates p < 0.001.

We subsequently visualized the expression and/or deposition of proteins associated with mesenchymal and endothelial cells in the native bone marrow. Initially, we aimed to distinguish arteriolar and sinusoidal endothelial cells by Sca-1 and podoplanin (PDPN) expression^50^. Though we did not observe co-staining of CD31^+^ endothelial cells with PDPN or Sca-1, we did observe strong staining of MSCs with PDPN (**Fig 2A**); further we observed no difference in overlap of CD31^+^ ECs with PDPN^+^ MSCs in normoxia or hypoxia. Networks exhibited laminin, an important extracellular matrix molecule enriched in the walls lining endothelial cells^51^; the levels of laminin at day 7 were still low but qualitatively higher in normoxia (**Fig 2B**). Lastly, networks stained for tight junction proteins Occludin (OCC) and ZO-1 showed an appreciable amount of ZO-1 regardless of oxygen tension, suggesting initial maturation of PVN networks in GelMA hydrogels (**Fig 2C**).

**Figure 2.**
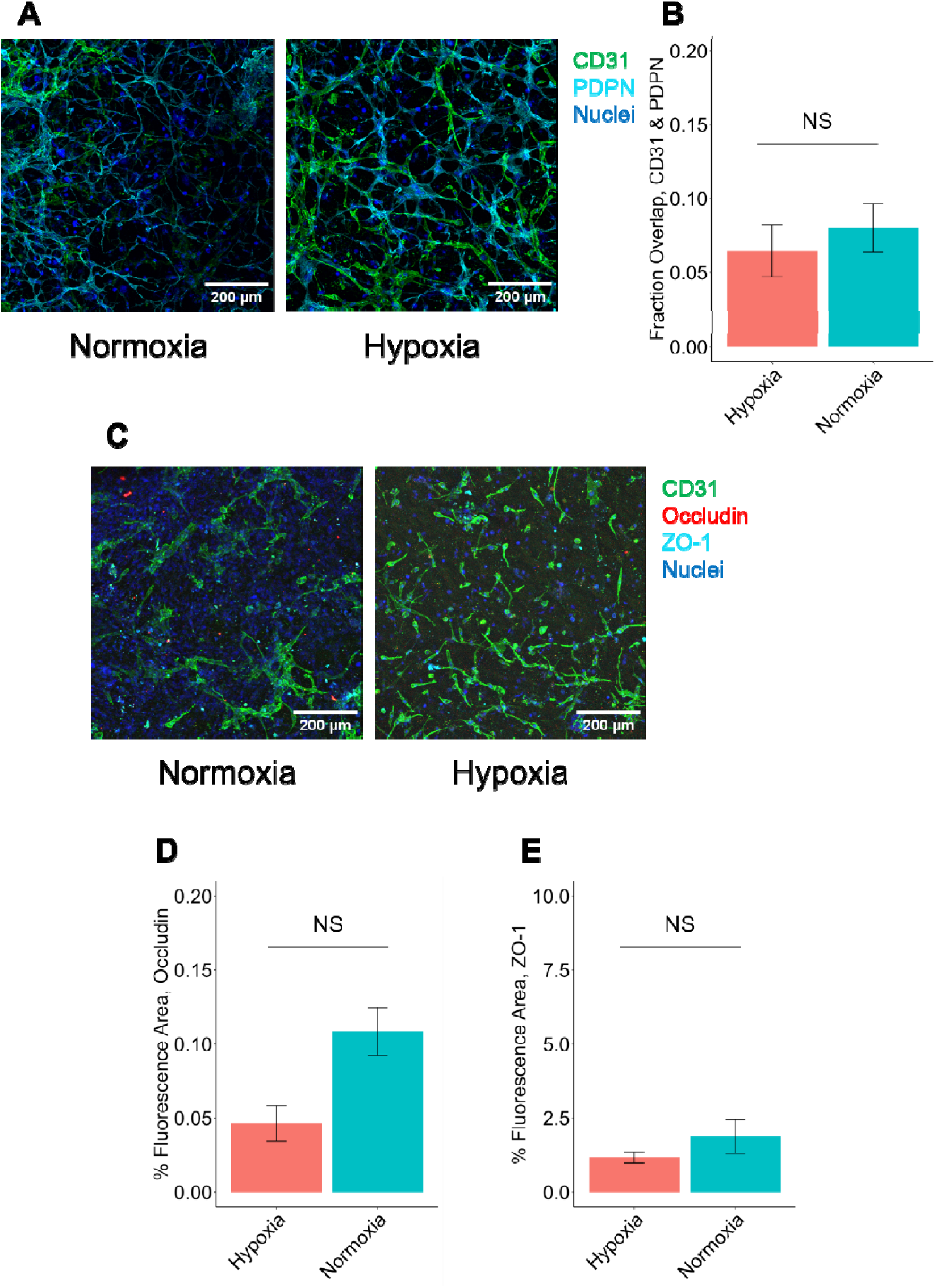
**A** Example images of PVN networks stained for CD31, Sca-1, PDPN, and Nuclei. **B** Fractional overlap between CD31 and PDPN is presented as a measure of colocalization. **C** Example images of PVN networks stained for CD31, Occludin, ZO-1 and Nuclei. Subsequent barplots quantify the deposition of **D** OCC and **E** ZO-1. For each condition, 3 images were taken to characterize each of N = 3 gels before analytical comparison. Barplots represent mean ± SEM.

### 3.2 Secretion of Angiogenesis- and Hematopoiesis-related Cytokines

Conditioned media from PVN hydrogels maintained in hypoxia contained many cytokines related to vasculature formation as well as HSC regulation (**Fig 3A-B**). Most factors were more highly expressed in normoxic vs. hypoxic cultures. CXCL12, a potent regulator of HSC maintenance and retention within the bone marrow, was more highly expressed under normoxic conditions. Interestingly, osteopontin (OPN), a factor known to promote HSC quiescence, was more highly expressed under hypoxic culture. Other factors more highly expressed in hypoxia include IGFBP-3, MCP-1, and MMP3. Because the differences observed between normoxia-cultured and hypoxia-cultured secretomes were relatively small, we proceeded with normoxia-cultured PVN medium in subsequent experiments.

**Figure 3.**
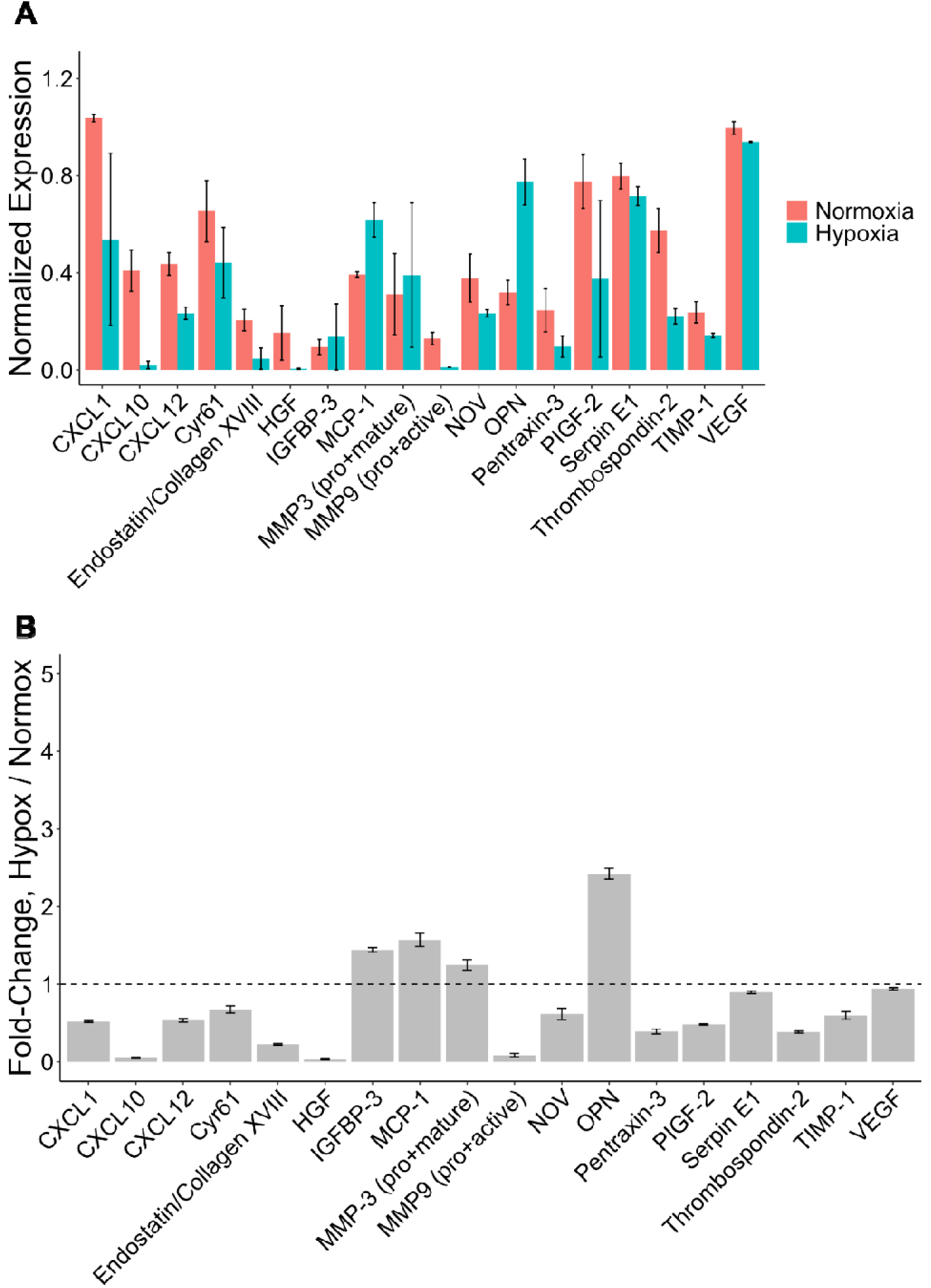
**A** Relative expression of detected cytokines secreted from BMECs and MSCs encapsulated in GelMA gels under normoxic and hypoxic conditions Data normalized to reference spots. Analysis performed on N = 2 batches of normoxic and N = 2 batches of hypoxic conditioned media with duplicate measurements taken on each membrane. Plots represent mean ± standard error of the mean. **B** Fold-change presented as: (Hypoxic cytokine level) / (Normoxic cytokine level).

### 3.3 Culture of HSPCs in Hypoxia with PVN Conditioned Media

Murine LSKs were then encapsulated in GelMA hydrogels and cultured in normoxia or hypoxia with or without conditioned media from the perivascular culture. After 4 days of culture, cells were analyzed via flow cytometry. The incorporation of PVN medium had a significant impact on cell expansion and the phenotypic composition of the HSPC compartment. When cultured with PVN medium, the total number of cells doubled for both normoxic and hypoxic cultures (**Fig 4A**). While hypoxic culture was generally less impactful than PVN medium, hypoxic culture induced a shift toward increased Lin^-^ (more primitive) cells (**Fig 4B**). Examining the relative expansion of the Lin^-^ vs. Lin^+^ cell fractions (day 4 vs. starting), both populations showed significant increases in response to PVN conditioned media (**Fig 4C-D**). Examining the HSPC fraction, we observed an even greater effect of the PVN medium. Here, we observed 2.70x and 3.19x fold-change increases in LSKs in normoxic and hypoxic cultures, respectively, in response to PVN medium (**Fig 4E**) as well as phenotypic shifts within the HSPC compartment (**Fig 4F-H**). While the fraction of LT-HSCs was unaffected, the fraction of ST-HSCs was reduced and the fraction of MPPs was increased. It is important to note that these effects on ST-HSCs and MPPs were driven by differential expansion rates of these populations rather than a loss of ST-HSCs (**Fig S3A-C**).

**Figure 4.**
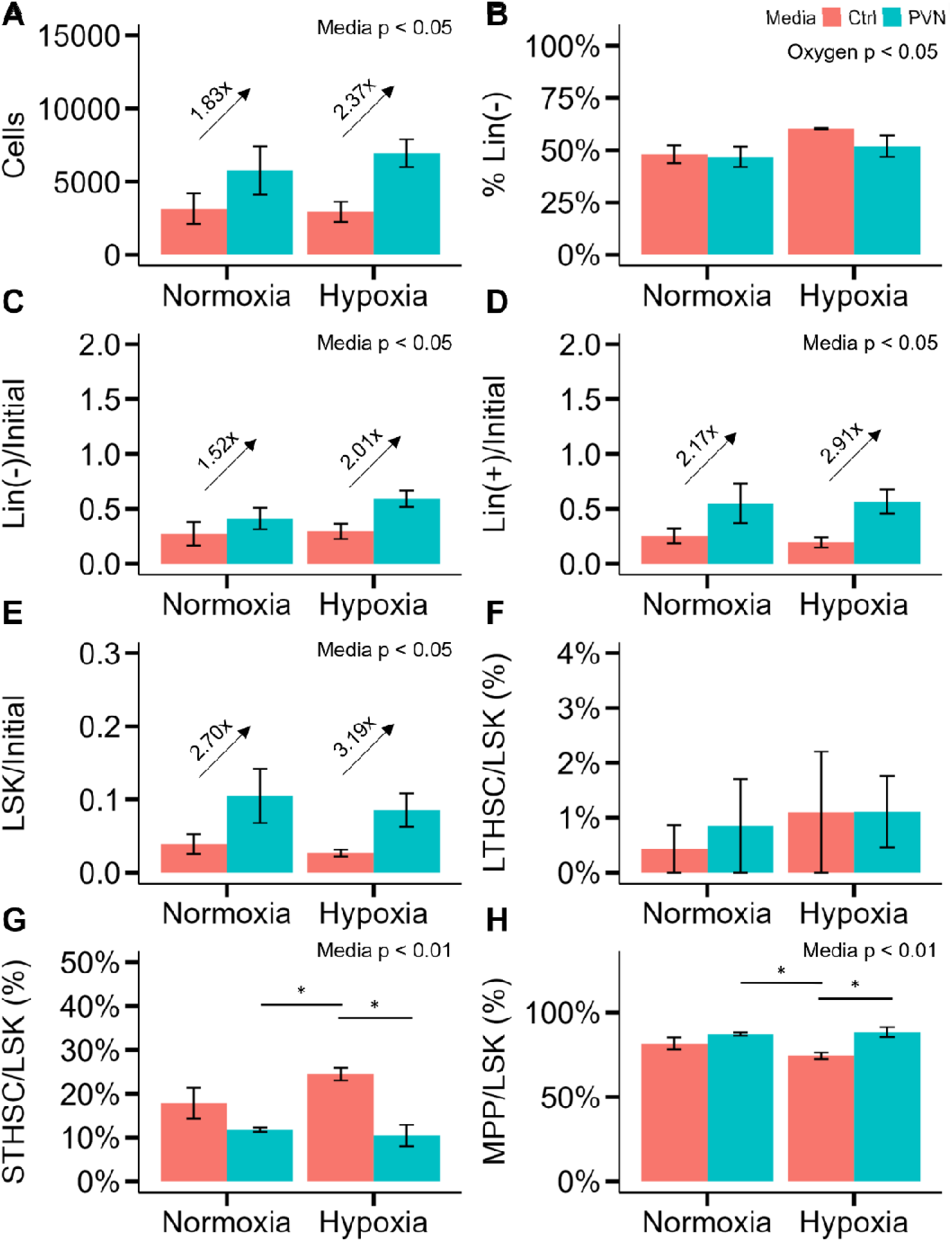
Barplots report mean ± SEM at day 4 of HSPC numbers and distributions as a function of oxygen tension and media as follows: **A** cells; **B** fraction of Lin^-^ cells, **C** Lin^-^ cells normalized to the starting cell number, **D** Lin^+^ cells normalized to the starting cell number, **E** LSKs normalized to the starting cell number, **F** LT-HSCs as a fraction of the LSK compartment, **G** ST-HSCs as a fraction of the LSK compartment, and **H** MPPs as a fraction of the LSK compartment. Initial number of cells was 6000. N = 5 gels for Hypoxia/Control condition; N = 4 gels for all other conditions. Significance of factors in Kruskal-Wallis or 2-way ANOVA is reported in the upper-right corner of each plot. * denotes p < 0.05 between individual groups.

### 3.4 Mitigation of Oxygen Shock and Culture of Hematopoietic Stem and Progenitor Cells in Hypoxia

After observing limited impact of hypoxia on HSPC phenotype, we hypothesized that EPHOSS experienced during HSPC isolation may mask the effect of hypoxic culture, since even brief oxygen exposure can alter HSPC phenotype and expansion. To test this hypothesis, we isolated HSPCs with and without CSA-loaded buffers, then performed GelMA cultures under normoxic and hypoxic conditions. Here, hypoxic culture significantly increased cell expansion (**Fig 5A**) particularly for HSPCs isolated with CSA to suppress oxygen shock. The balance of Lin^-^ and Lin^+^ cells at the end of culture was shifted toward Lin^-^ (**Fig 5B**) when cells were cultured in hypoxia – again, with a larger effect in cells isolated with CSA. The expansion of Lin^-^ cells was greatly increased with hypoxic culture for HSPCs that were isolated with CSA (**Fig 5C**). CSA isolation also slightly elevated Lin^+^ expansion under hypoxia (**Fig 5D**).

**Figure 5.**
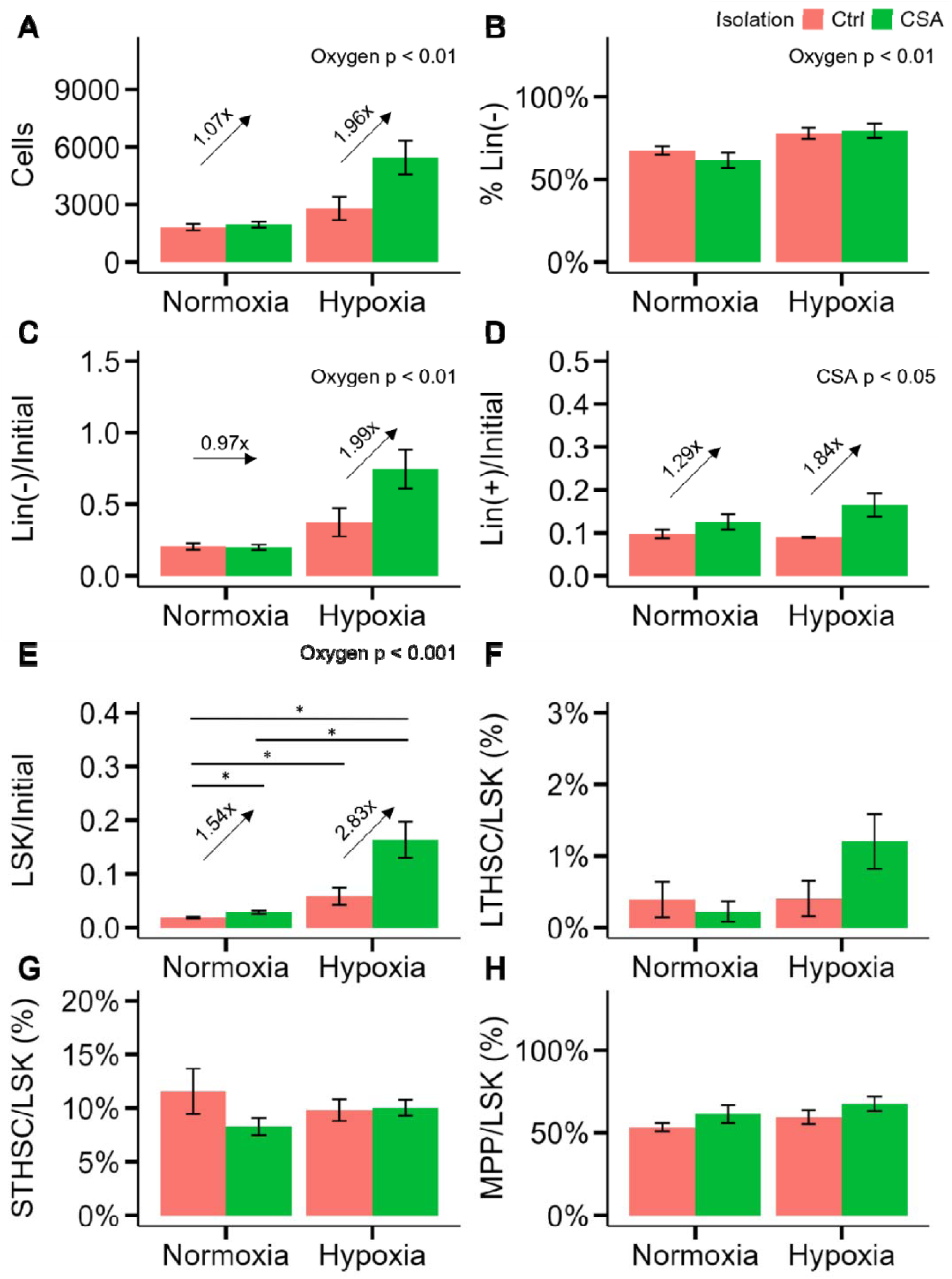
Barplots report mean ± SEM at day 4 of HSPCs numbers and distributions as a function of oxygen tension and isolation buffer as follows: **A** cells; **B** fraction of Lin^-^ cells, **C** Lin^-^ cells normalized to the starting cell number, **D** Lin^+^ cells normalized to the starting cell number, **E** LSKs normalized to the starting cell number, **F** LT-HSCs as a fraction of the LSK compartment, **G** ST-HSCs as a fraction of the LSK compartment, and **H** MPPs as a fraction of the LSK compartment. Initial number of cells was 6000. N = 5 gels for all conditions. Significance of factors in Kruskal-Wallis or 2-way ANOVA is reported in the upper-right corner of each plot. * denotes p < 0.05 between individual groups.

Excitingly, CSA isolation paired with hypoxic culture induced significant increases in the number of LSKs during culture (**Fig 5E**). As the fractional composition of the HSPC compartment was unaffected (**Fig 5F-H**), the increase in LSKs after CSA isolation and hypoxic culture was associated with increases in absolute numbers of LT-HSCs, ST-HSCs, and MPPs (**Fig S4A-C**).

### 3.5 Coordinated Impact of Hypoxia and Perivascular Niche Conditioned Medium with CSA Isolation

Having demonstrated the importance of mitigating EPHOSS during HSC isolation, we examined the effect of hypoxia and PVN conditioned medium on HSPCs isolated with CSA to reduce EPHOSS. Here, PVN conditioned medium induced a significant expansion of cells (**Fig 6A**). Hypoxic culture shifted the fractional composition of cells toward the Lin^-^ phenotype (**Fig 6B**, **Fig 6C**), while incorporation of the PVN conditioned medium greatly increased expansion of Lin^+^ cells (**Fig 6D**). We also observed a similar effect of hypoxia on CSA-isolated HSPCs, with an increase (p = 0.088) in the number of LSKs for CSA-isolated, hypoxia-cultured cells versus those cultured in normoxia (**Fig 6E**).

**Figure 6.**
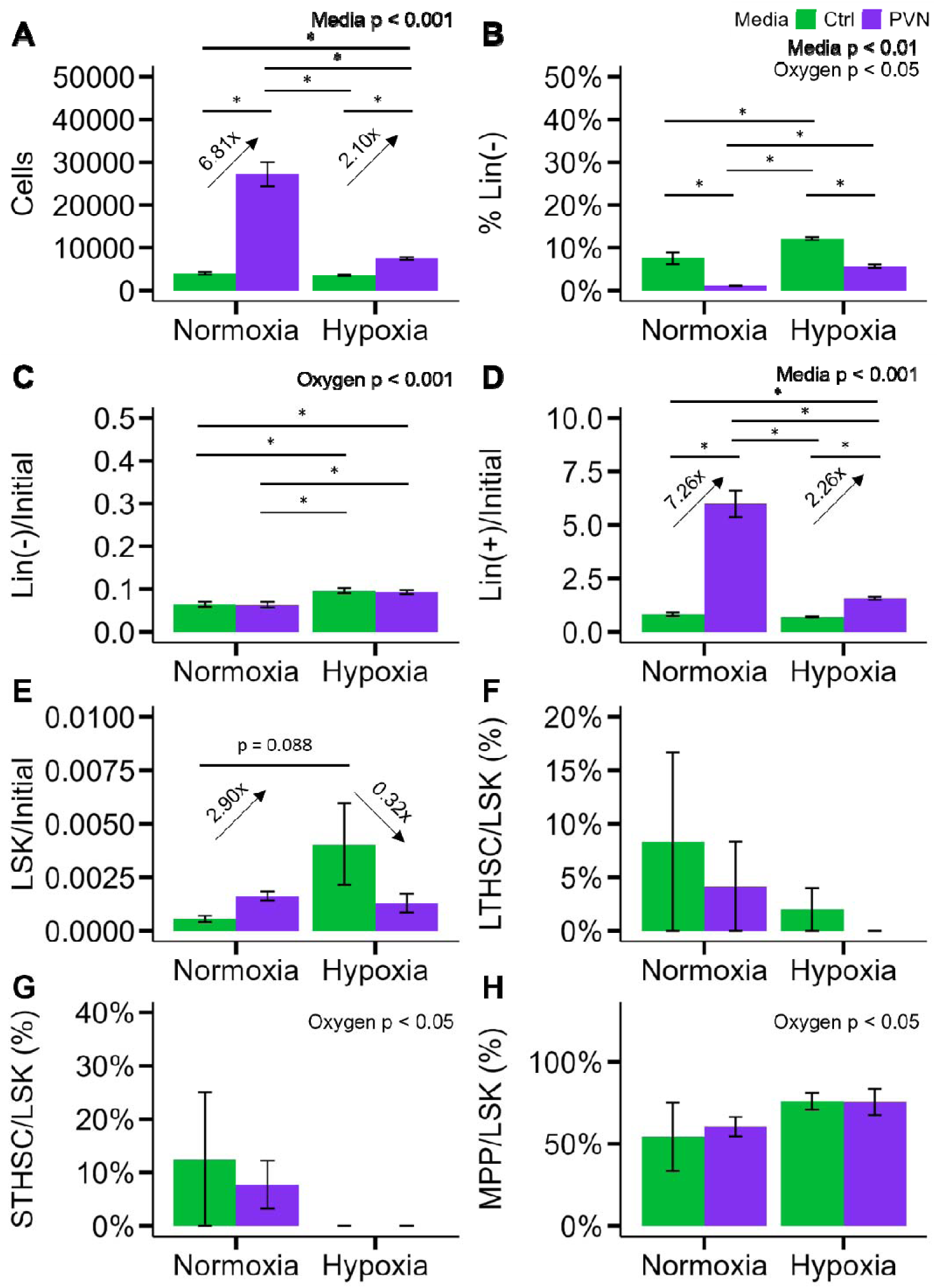
Barplots report mean ± SEM at day 4 of HSPCs numbers and distributions as a function of oxygen tension and media after isolation with CSA-containing buffer: **A** cells; **B** fraction of Lin^-^ cells, **C** Lin^-^ cells normalized to the starting cell number, **D** Lin^+^ cells normalized to the starting cell number, **E** LSKs normalized to the starting cell number, **F** LT-HSCs as a fraction of the LSK compartment, **G** ST-HSCs as a fraction of the LSK compartment, and **H** MPPs as a fraction of the LSK compartment. Initial number of cells was 4500. N = 5 gels for hypoxia-cultured gels; N = 4 gels Normoxia-cultured gels. Significance of factors in Kruskal-Wallis or 2-way ANOVA is reported in the upper-right corner of each plot. * denotes p < 0.05 between individual groups.

## 4. Discussion

In this study, we examined the effects of physiologically relevant oxygen levels on the formation of bone marrow vasculature and the subsequent effect of hypoxia and PVN conditioned media on murine HSPC expansion. We find that PVN conditioned media provide a powerful signal to influence HSPC expansion and favorable differentiation. Notably, we observe the effect of hypoxic culture on HSPCs is largely obscured by HSPC isolation conditions, and that isolation using methods to reduce oxygen-associated EPHOSS is important for in vitro studies of HSC fate.

We first examined the formation of 3D PVN networks in gelatin hydrogels using murine BMECs and MSCs. Quantitative comparison of network formation under hypoxia vs. normoxia shows a lower total network length, vessel density, and branch point density in hypoxic cultures. While hypoxia can induce the upregulation of angiogenesis-related genes, the effect was not seen in short term hydrogel culture. Ongoing work is examining expression profiles of a larger slate of angiogenic genes (VEGF, Angiopoietin-1, Tie2, PDGFβ) as well as the possibility of adding exogenous factors to further enhance angiogenic activity. However, closer inspection of the literature reveals a study by Gawlitta et al. attempting to prevascularize bone constructs before implantation under low oxygen conditions (5%) that similarly showed reduced network formation^52^. Here, pellets of endothelial colony forming cells and multipotent stromal cells failed to form prevascular structures under hypoxia, but showed robust formation under normoxia. They hypothesized that while hypoxia is a strong inducer of angiogenesis (formation of new blood vessels from existing vessels), it may not provide sufficient metabolic support to drive effective vasculogenesis (de novo formation of a vascular network). They also hypothesized that due to the large size of the clusters and diffusion limits, cells in the center of normoxic constructs may experience hypoxia, while those in hypoxic conditions would experience even lower oxygen levels. This work differs from our study because they relied upon 3D clusters of cells rather than scaffolds or hydrogels; however, diffusive transport within three-dimensional hydrogels may also further reduce oxygen and small molecule availability in our PVN model^30,41^, especially as cells proliferate to high density and consume more oxygen. Metabolic activity at day 7 of our perivascular cultures shows cells maintained under hypoxia had significantly lower metabolic activity, which may be due to a lower number of cells or an equal number of less active cells. Further study will be needed to more accurately determine which is the case, such as via quantification of the total number of cells at day 7 in each culture or examination of the proliferative activity of cells within the PVN model via immunofluorescent analysis of Ki67 or Edu activity^53,54^.

The strong signal of PDPN on MSCs in this study will be useful for network visualization in future studies. PDPN is primarily expressed on certain types of mesenchymal stromal cells^55^, and is critical for the development and function of several tissues, structures including lung, heart, and lymphatic tissues. This protein has also been studied as a cancer biomarker and as a signifier of an increase in the epithelial mesenchymal transition associated with motility and metastasis^56^. PDPN interacts with CLEC-2, a receptor expressed on platelets and some immune cells, to induce platelet aggregation and lymphatic vessel development^55,57^. Though not specifically evaluated here, platelet and immune cell numbers and function could be investigated in relation to the presence of these PDPN^+^ MSCs in future more complex marrow models. Analysis of conditioned media using an angiogenesis cytokine array revealed expression of a wide array of factors by co-cultured BMECs and MSCs maintained in either hypoxia or normoxia. Normoxic cultures had higher expression of matrix remodeling related cytokines, including MMP9, which likely contributed to the greater network formation. Normoxic PVN cultures also showed higher expression of CXCL12, an HSC niche factor known to play a significant role in HSC retention and maintenance^58^. Hypoxic PVN cultures presented a higher level of OPN, a negative regulator of HSC pool size, and is thereby associated with a higher degree of quiescence^59,60^.

The importance of oxygen tension is generally accepted, but demonstrating this importance proved less straightforward than simply culturing cells at varying levels of oxygen. All experiments presented a phenotypic shift toward Lin^-^ hematopoietic progenitor cells with hypoxic culture, but other effects were dependent upon HSPC isolation using CSA to mitigate the impact of oxygen shock. Phenotypic and functional shifts in cultured HSPCs were recently observed in 2D culture without the use of CSA to mitigate oxygen shock^28^, but these phenotypic changes were only observed from day 14 onward and functional capacity was evaluated after 28 days. Though CSA-based isolation may not be necessary to observe the eventual effects of hypoxia, it stands to reason that CSA-based isolation may enhance or accelerate expansion in hypoxic culture by ameliorating oxygen shock and bypassing this phenotypic recovery. HSPCs cultured in both hypoxia and normoxia showed improved retention of a subset of primitive LSKs over the 4-day culture period in the presence of PVN conditioned media (**Fig 4E**), but this effect was diminished for 4-day cultures when cells were isolated with CSA and cultured in hypoxia (**Fig 6E**).

Overall, these studies suggest oxygen tension is a significant variable not just in the formation of biomaterial models of the bone marrow, but in the isolation of cells incorporated into the models. Though the PVN medium applied had similar effects on the expansion of Lin^+^ cells, its effects on Lin^-^ expansion and the HSPC compartment were more variable. Previous work by Eliasson et al. culturing HSPCs under 1% oxygen showed an increase in the proportion of quiescent HSCs, but this concurrently reduced expansion of hematopoietic cells^61^. Importantly, we observed increases in the numbers of phenotypic HSPCs as well as Lin^+^ hematopoietic cells, suggesting that additional factors beyond exogenous supplementation, such as biomaterials, metabolic conditions, and co-cultured cells will enhance methods for HSC expansion in future extensions of this work. Here, we observed the presence of CXCL12 and OPN, but other secreted factors have been implicated in HSC regulation, and may be tuned through co-culture conditions. Further studies are expected to reveal the precise phenotypes (i.e., arteriolar vs. sinusoidal) of hydrogel-based PVN cultures and explore direct co-culture of perivascular cells with HSPCs.

## 5. Conclusion

In this work, we evaluated the impact of hypoxic culture upon a GelMA hydrogel model of the bone marrow. We further explored the combined impact of mitigating EPHOSS in HSPC isolation, hypoxic culture, and conditioned media supplementation upon HSPC fate in this hydrogel model. PVN networks exhibited signs of preliminary maturation through expression of laminin, OCC, and ZO-1 under both normoxic and hypoxic conditions, and these culture conditions induced expression of key regulators of HSC maintenance and quiescence including CXCL12 and OPN. PVN conditioned medium showed the potential to enhance biomaterial-based expansion of HSPCs. Hypoxic culture consistently shifted the distribution of cells toward a more primitive Lin^-^ phenotype, and mitigation of EHPOSS with CSA significantly enhanced maintenance of HSPCs when followed by a hypoxic culture condition. These findings demonstrate the importance of isolation methods and hypoxic culture in engineered models of the bone marrow, introducing a key element to endeavors aimed at HSC maintenance and expansion with biomaterial platforms.

## Acknowledgements

Funding sources include the National Institute of Diabetes and Digestive and Kidney Diseases of the National Institutes of Health under Award Number 2 R01 DK099528, the National Institute of Dental and Craniofacial Research of the National Institutes of Health under Award Number R01 DE030491, and the National Cancer Institutes of the National Institutes of Health under Award Number R01 CA256481. The content is solely the responsibility of the authors and does not necessarily represent the official views of the NIH. The authors are also grateful for additional funding provided by the Department of Chemical & Biomolecular Engineering, Department of Bioengineering, the Carl R. Woese Institute for Genomic Biology, the Cancer Center at Illinois, and the Illinois Scholars Undergraduate Research Program at the University of Illinois Urbana-Champaign. The authors wish to thank the Roy J. Carver Biotechnology center, Cytometry and Microscopy to Omics Facility for providing expertise in Cytometry/Sorting, Microscopy and Omics analysis for our project. Specifically, we would like to thank Dr. Mayandi Sivaguru and Dr. Marcin Wozniak for their assistance with cell sorting. The authors also wish to thank the Core Facilities at the Carl R. Woese Institute for Genomic Biology for use of their LSM710 and LSM880 microscopes. In particular, we would like to thank Dr. Austin J. Cyphersmith and Dr. Duncan Nall for training and imaging assistance.

## Contributions (CRediT: Contributor Roles Taxonomy^62,63^)

**G.B. Thompson:** Conceptualization, Data curation, Formal Analysis, Visualization, Investigation, Methodology, Writing – original draft, Writing – review & editing. **V.R. Barnhouse:** Conceptualization, Data curation, Formal Analysis, Visualization, Investigation, Methodology. **S.K. Bierman:** Investigation, Methodology, **B.A.C. Harley:** Conceptualization, Resources, Project administration, Funding acquisition, Supervision, Writing – review & editing.

## Data Availability Statement

The data collected and used to support the findings of this study will be made available by the corresponding author, B.A.C. Harley, upon reasonable request.

