## Supplemental Information for "Influence of Hypoxia on a Biomaterial Model of the Bone Marrow Perivascular Niche"

^1^Dept. Chemical and Biomolecular Engineering

^2^Dept. Bioengineering

^3^Cancer Center at Illinois

^4^Carl R. Woese Institute for Genomic Biology

University of Illinois at Urbana-Champaign

Urbana, IL 61801

**Corresponding Author:**

B.A.C. Harley

Dept. of Chemical and Biomolecular Engineering

Cancer Center at Illinois

Carl R. Woese Institute for Genomic Biology

University of Illinois at Urbana-Champaign

110 Roger Adams Laboratory

600 S. Mathews Ave.

Urbana, IL 61801

**Table S1** Primary and secondary antibodies, sources, species, and concentrations for immunostaining of networks for morphology and protein deposition.

| **Primary vs. Secondary** | **Antibody** | **Vendor and Catalog Number** | **Host Species** | **Working concentration or dilution** |
| --- | --- | --- | --- | --- |
| Primary | Anti-CD31 | Abcam  ab7388 | Rat | 10 µg/mL |
| Primary | Anti-Sca-1 | Thermo Fisher  701919 | Rabbit | 2 µg/mL |
| Primary | Anti-PDPN | R&D Systems  AF3244 | Goat | 2 µg/mL |
| Primary | Anti-Laminin | Thermo Fisher  PA5-36300 | Rabbit | 10 µg/mL |
| Primary | Anti-Occludin | Thermo Fisher  710192 | Rabbit | 1 µg/mL |
| Primary | Anti-ZO-1 | Thermo Fisher  PA5-19090 | Goat | 10 µg/mL |
| Secondary | Donkey Anti-rat IgG AF 555 | Thermo Fisher A78945 | Donkey | 1:500 |
| Secondary | Donkey Anti-rabbit IgG H&L (AF 647) | Thermo Fisher A31573 | Donkey | 1:500 |
| Secondary | Donkey Anti-Goat IgG H&L (AF 488) | Thermo Fisher A11055 | Donkey | 1:500 |

**Table S2** Antibodies, sources, and concentrations used for sorting pre-enriched hematopoietic cells.

| **Antibody** | **Fluorophore** | **Vendor and Catalog Number** | **µL antibody / µL cell solution** |
| --- | --- | --- | --- |
| CD5 | FITC | Thermo Fisher 11-0051-82 | 1:100 |
| B220 | FITC | Thermo Fisher 11-0452-82 | 1:100 |
| CD8a | FITC | Thermo Fisher 11-0081-82 | 1:100 |
| Gr-1 | FITC | Thermo Fisher 11-5931-82 | 0.25:100 |
| Ter-119 | FITC | Thermo Fisher 11-5921-82 | 0.5:100 |
| CD11b | FITC | Thermo Fisher 11-0112-82 | 1:100 |
| Sca-1 | PE | Thermo Fisher 12-5981-83 | 1.25:100 |
| c-Kit | APC-780 | Thermo Fisher 47-1172-82 | 0.625:100 |

**Table S3** Antibodies, sources, and concentrations used for endpoint flow cytometry analysis of hematopoietic populations.

| **Antibody** | **Fluorophore** | **Vendor and Catalog Number** | **µL antibody / µL cell solution** |
| --- | --- | --- | --- |
| Mouse Lineage Antibody Cocktail | PerCP-Cy5.5 | BD Biosciences  561317 | 20:100 |
| Sca-1 | PE-Cy7 | BD Biosciences  561021 | 0.625:100 |
| c-Kit | R718 | BD Biosciences  567841 | 1.25:100 |
| CD150 | PE | BD Biosciences  562651 | 2.5:100 |
| CD48 | APC | BD Biosciences  562746 | 0.6:100 |
| N/A | DAPI (1 mg/mL) | BD Biosciences  564907 | 5:300 |

**Table S4** Summary of network morphological characteristics presented in Figure 1. Values represent mean ± SEM.

| **Oxygen** | **Network Length (mm)** | **Branch points/mm^3^** | **Vessels/mm^3^** | **Branch Length (mm)** |
| --- | --- | --- | --- | --- |
| 18% | 98.1 ± 22.6 | 1800.9 ± 538.4 | 4256.0 ± 1008.4 | 0.068 ± 0.001 |
| 1.5% | 55.6 ± 12.6 | 717.5 ± 243.5 | 2570.5 ± 584.7 | 0.064 ± 0.003 |

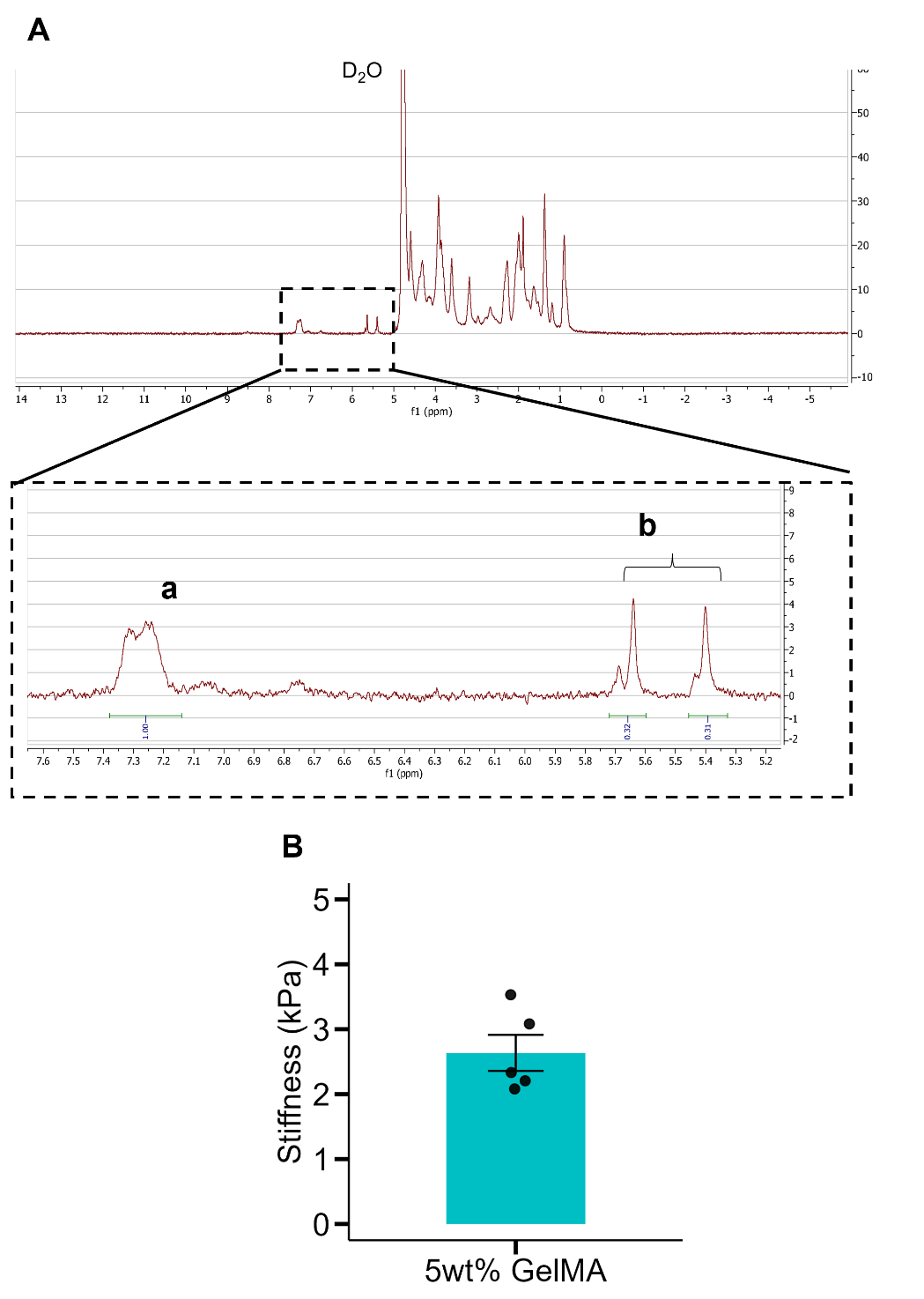

**Figure S1 A** NMR spectrum for GelMA. Using phenylalanine peak (a) as a reference, the relative degree of functionalization is 0.63 by integrating and summing the peaks associated with methacrylamide functionalization (b). **B** Stiffness measured via unconfined compression of GelMA hydrogels. N = 5.

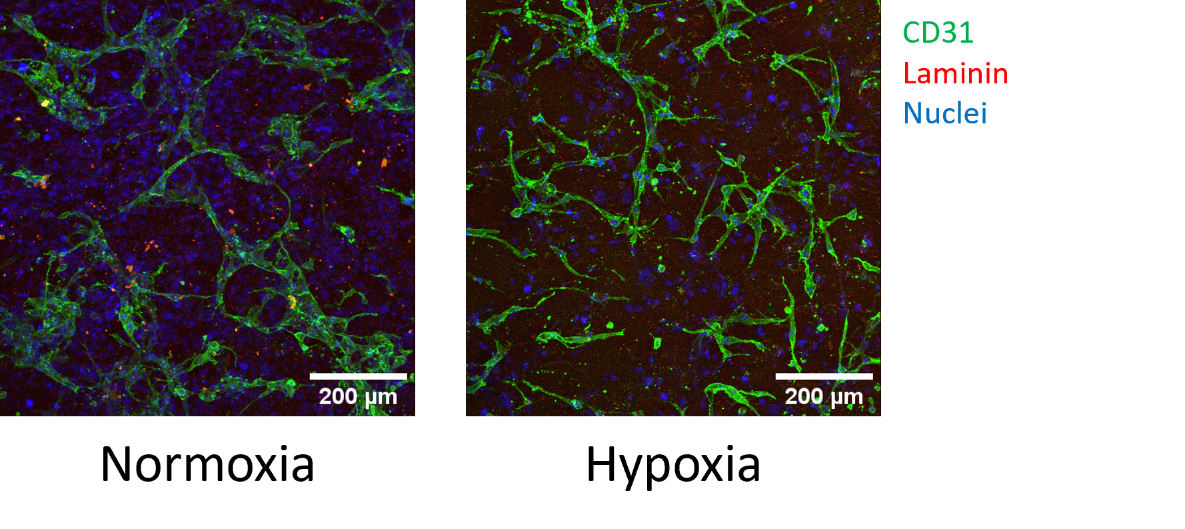

**Figure S2** Example images of endothelial networks stained for CD31 and laminin.

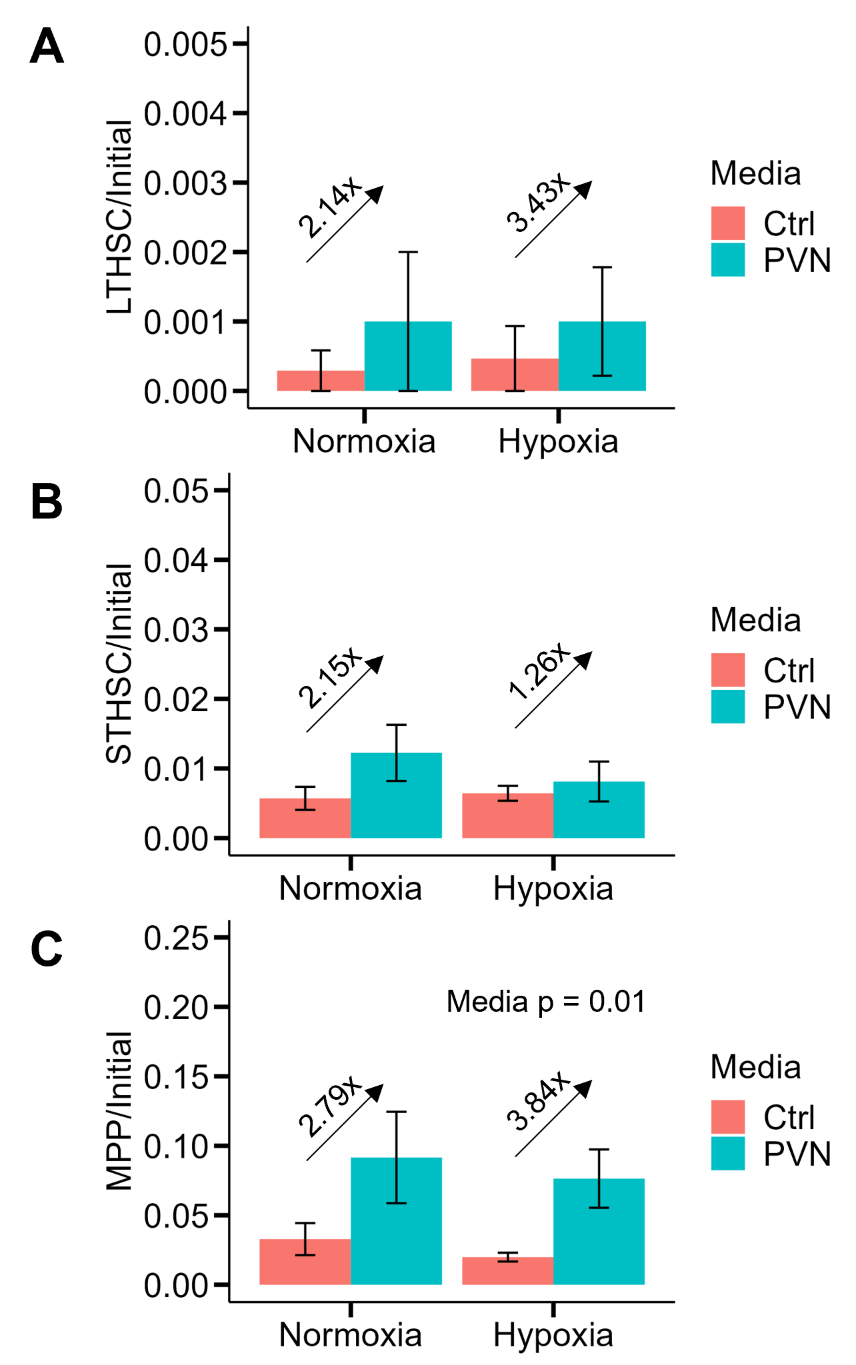

**Figure S3** Barplots report mean ± SEM at day 4 of HSPC numbers as a function of oxygen tension and media supplementation, normalized to the initial cell count as follows: **A** LT-HSC, **B** ST-HSC, and **C** MPP.

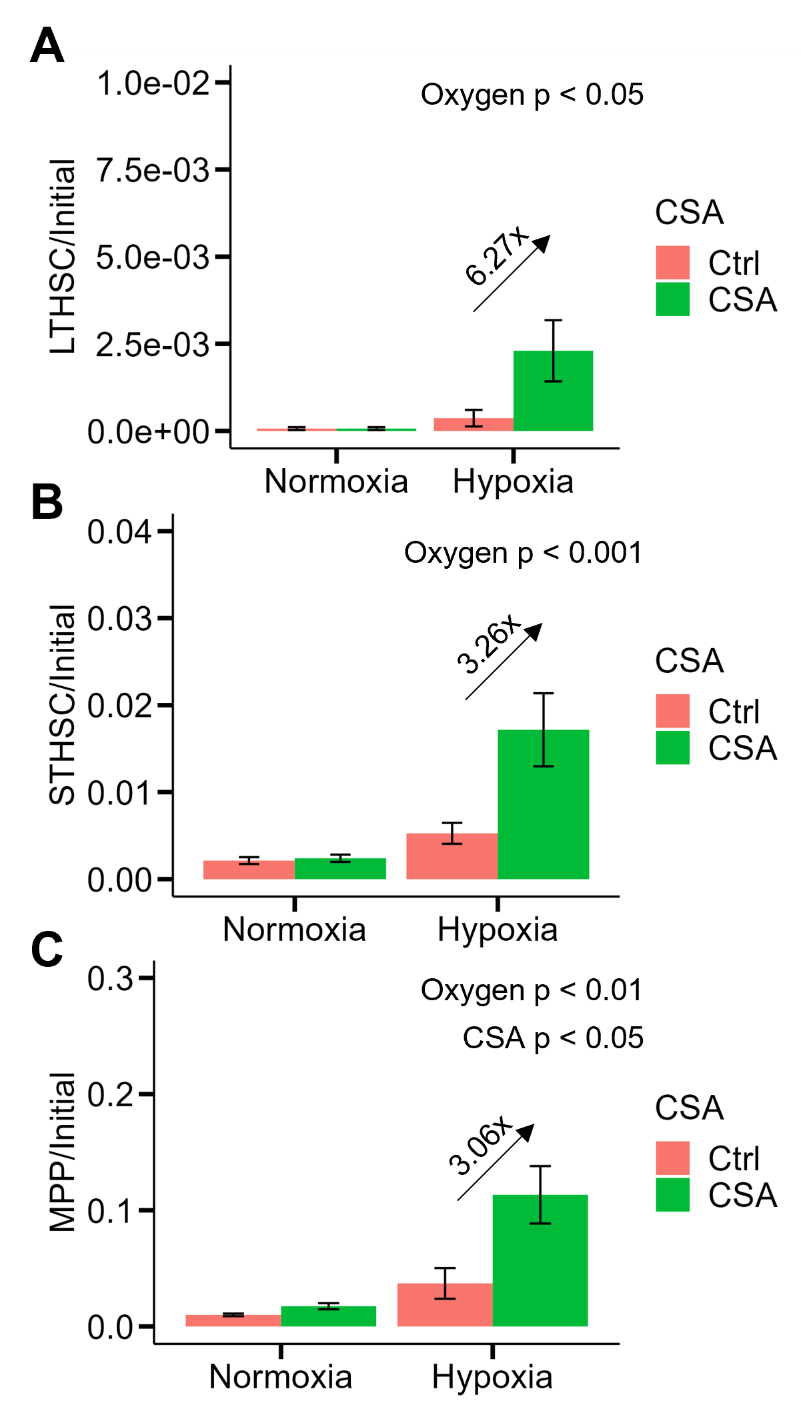

**Figure S4** Barplots report mean ± SEM at day 4 of HSPC numbers as a function of oxygen tension and isolation method, normalized to the initial cell count as follows: **A** LT-HSC, **B** ST-HSC, and **C** MPP.

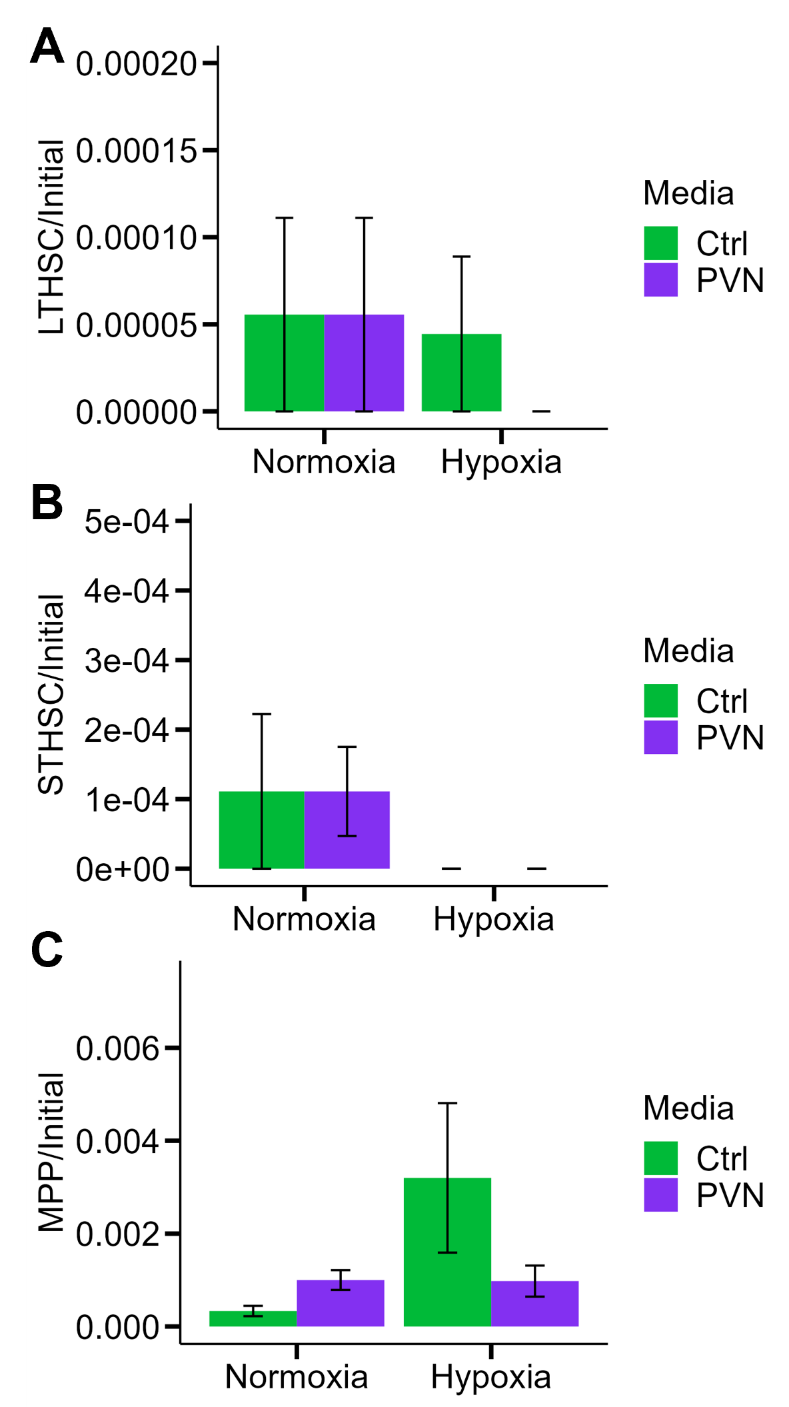

**Figure S5** Barplots report mean ± SEM at day 4 of HSPC numbers as a function of oxygen tension and media supplementation, normalized to the initial cell count as follows: **A** LT-HSC, **B** ST-HSC, and **C** MPP. All cells were isolated using buffers that contained CSA.
